# Activation of somatostatin interneurons by nicotinic modulator Lypd6 enhances plasticity and functional recovery in the adult mouse visual cortex

**DOI:** 10.1101/155465

**Authors:** Masato Sadahiro, Michael P. Demars, Poromendro Burman, Priscilla Yevoo, Andreas Zimmer, Hirofumi Morishita

## Abstract

The limitation of plasticity in the adult brain impedes functional recovery later in life from brain injury or disease. This pressing clinical issue may be resolved by enhancing plasticity in the adult brain. One strategy for triggering robust plasticity in adulthood is to reproduce one of the hallmark physiological events of experience-dependent plasticity observed during the juvenile critical period – rapidly reduce the activity of parvalbumin (PV)-expressing interneurons and disinhibit local excitatory neurons. This may be achieved through enhancement of local inhibitory inputs, particularly those of somatostatin (SST)-expressing interneurons. However, to date the means for manipulating SST interneurons for enhancing cortical plasticity in the adult brain are not known. We show that SST interneuron-selective overexpression of Lypd6, an endogenous nicotinic signaling modulator, enhances ocular dominance plasticity in the adult primary visual cortex (V1). Lypd6 overexpression mediates a rapid experience-dependent increase in the visually evoked activity of SST interneurons as well as a simultaneous reduction in PV interneuron activity and disinhibition of excitatory neurons. Recapitulating this transient activation of SST interneurons using chemogenetics similarly enhanced V1 plasticity. Notably, we show that SST-selective Lypd6 overexpression restores visual acuity in amblyopic mice that underwent early long-term monocular deprivation. Our data in both male and female mice reveal selective modulation of SST interneurons and a putative downstream circuit mechanism as an effective method for enhancing experience-dependent cortical plasticity as well as functional recovery in adulthood.

**Significance Statement:** The decline of cortical plasticity after closure of juvenile critical period consolidates neural circuits and behavior, but this limits functional recovery from brain diseases and dysfunctions in later life. Here we show that activation of cortical SST interneurons by Lypd6, an endogenous modulator of nicotinic acetylcholine receptors (nAChRs), enhances experience-dependent plasticity and recovery from amblyopia in adulthood. This manipulation triggers rapid reduction of PV interneuron activity and disinhibition of excitatory neurons, which are known hallmarks of cortical plasticity during juvenile critical periods. Our study demonstrates modulation of SST interneurons by Lypd6 to achieve robust levels of cortical plasticity in the adult brain and may provide promising targets for restoring brain function in the event of brain trauma or disease.

## Introduction

Experience-dependent plasticity heightens during juvenile critical periods to dynamically shape brain function and behavior, then declines into adulthood to ensure stability. This poses a challenge for functional recovery in later life after brain injury or disease (Hensch, 2004; Knudsen, 2004). For instance, incongruous vision during early childhood results in amblyopia, a vision disorder marked by an enduring loss of visual acuity with limited treatment options in adulthood (Wiesel, 1982). Restoration of heightened plasticity in the adult brain is thought to support restoration of function from such debilitating conditions (Hensch and Quinlan, 2018).

Ocular dominance (OD) plasticity, wherein a brief monocular visual deprivation in early life results in a robust shift of the responsiveness of the primary visual cortex (V1) from the deprived eye to the non-deprived eye, has been a long-standing model for understanding the fundamental regulatory mechanisms of critical periods and discovering novel targets for restoring heightened cortical plasticity in adulthood. Recent studies identified one of key hallmarks of V1 plasticity during the critical period where a transient disinhibition of the excitatory cortical network occurs immediately following visual deprivation to rapidly restore cortical activity to levels before visual deprivation to initiate competitive plasticity. This disinhibition is mediated by a reduction of perisomatic inhibition by the parvalbumin (PV)-expressing GABAergic interneurons (Hengen et al., 2013; Kuhlman et al., 2013; Feese et al., 2018). Importantly, this rapid deprivation induced reduction of PV interneuron activity and disinhibition of local excitatory neurons does not occur in adulthood (Kuhlman et al., 2013). Notably, artificially replicating this transient reduction in PV interneuron activity in adulthood through inhibitory chemogenetics robustly reactivates the juvenile form of OD plasticity marked by depression of deprived eye response (Kuhlman et al., 2013). This indicates that disinhibition of the excitatory network should be a fundamental aim for identifying molecular or circuit targets for enhancing experience-dependent cortical plasticity and developing therapeutic strategies for recovery from brain disorders such as amblyopia.

Controlled inhibition of cortical PV interneurons may potentially be achieved through enhancement of local inhibitory inputs, particularly from somatostatin (SST)-expressing GABAergic interneurons. SST interneurons inhibit PV interneurons twice as potently as they do local pyramidal excitatory neurons (Cottam et al., 2013), and the SST-PV disinhibitory microcircuit is well-recognized across cortical functions (Hu et al., 2011; Hioki et al., 2013; Pfeffer et al., 2013; Xu et al., 2013; Kato et al., 2015; Sturgill and Isaacson, 2015; Xu et al., 2019; Yaeger et al., 2019). However, only until recently the potential for of SST interneurons in reactivating V1 plasticity in the adult brain remained underexplored (Tang et al., 2014; Davis et al., 2015; Fu et al., 2015). Neuromodulation of SST interneurons, particularly through cholinergic signaling (Jia et al., 2010; Leao et al., 2012; Zhao-Shea et al., 2013; Chen et al., 2015), may represent one potential mean for enhancing plasticity in the adult brain. A recent study revealed that V1 SST interneurons during the critical period have heightened sensitivity to cholinergic modulation, and with visual activity this mediates fundamental mechanisms of cortical plasticity, including somatic disinhibition as well as branch-specific dendritic inhibition of pyramidal neurons (Yaeger et al., 2019). While this sensitivity declines in adulthood, the said study found that optogenetic stimulation of SST interneurons, concurrent with arousal, reactivates somatic disinhibition and branch-specific dendritic inhibition. While there are no known molecular targets in SST interneurons in the adult brain for enhancing cortical plasticity yet, perhaps selective enhancement of cholinergic modulation in V1 SST interneurons may be one strategy.

Our recent study revealed that Lypd6, an endogenous modulator of nicotinic acetylcholine receptors (nAChRs), is highly enriched in V1 SST interneurons (Demars and Morishita, 2014). The Ly6 family of proteins that includes Lypd6 are GPI anchored to the membrane, share unique toxin-like structures, and are known to bind to the extracellular face of nAChRs and cell-autonomously regulate their signaling (Miwa et al., 2012). Lypd6 is also known to potentiate calcium currents through nAChRs (Darvas et al., 2009). In this study, we sought to elucidate the potential for targeting nicotinic modulation through Lypd6 in SST interneurons in enhancing plasticity and functional recovery in adulthood.

## Materials and Methods

### Animals

All mice were housed in groups of 2–5 together with littermates of the same sex in standard and uniform cage sizes (199 mm × 391 mm × 160 mm: width × depth × height, GM500, Tecniplast) under a 12hr light:dark cycle (lights on at 7:00AM: lights off at 7:00PM) in a facility with constant temperature (23°C) and ad libitum access to food and water. Wild-type C57Bl/6 mice were obtained from Jackson laboratory and Charles River. Both male and female were used. Somatostatin-IRES-Cre (SST-Cre: Jackson laboratory #013044) was purchased and bred in-house. Bigenic lines were created through targeted breeding of above strains. Lypd6Tg mice were originally generated by A.Z.(Darvas et al., 2009), and transferred to H.M. and back-crossed to C57Bl/6. All animal protocols were approved by the Institutional Care and Use Committee (IACUC) at Icahn School of Medicine at Mount Sinai.

### Monocular deprivation

Monocular deprivation procedure for animals that underwent either 4 days (4d MD), 1 day (1d MD), or long-term (LTMD) monocular deprivation was conducted on the contralateral eye and with ipsilateral eye left open. The mice were anesthetized with isoflurane during the entire procedure. Eyelid margins were trimmed using an iris scissor and one eye was sutured closed for one or four days. Following MD, mice were returned to their home cage prior to extracellular recording and subsequent euthanasia. For both ocular dominance plasticity measurements (4DMD) and cell type specific visually evoked firing rate assessments (1DMD), the deprived eye was reopened just prior to the start of recording. For amblyopic mice (LTMD), the deprived eye was opened after the closure of V1 critical period (P33) and maintained to stay open until the day of visual evoked potential recordings (P60).

### Generation and validation of AAV-Lypd6

*Lypd6* was amplified from a cDNA library derived from whole mouse V1, subcloned into a pcDNA3.1 (-) vector. Subclonings were performed using an isothermal DNA assembly method (Gibson Assembly; New England Biolabs) and transformed into *E. Coli*. Colonies with correct insert were identified through DNA sequencing (Genewiz), cultured and then isolated using a Hi-speed Midiprep kit (Qiagen). Expression was first examined by transfection of N2A cells *in vitro*. To create the pAAV vector, an inverted bicistronic 2A sequence was inserted into pAAV-Ef1α-DIO-EGFP-WPRE-pA (Addgene#37084) upstream of EGFP by PCR linearization and overhang production on pAAV vector. The pcDNA3.1(-)-Lypd6 vector was used as a template for the Lypd6 insert which was subsequently inserted into the pAAV-DIO-EGFP-2A vector as described above to create a pAAV-DIO-EGFP-2A-Lypd6-WPRE-pA vector. After sequence verification, a large culture and Maxiprep isolation produced a purified vector that was sent to the UNC viral core for viral packaging using an AAV8 serotype. To confirm viral efficiency, AAVs were stereotaxically injected (see below for stereotaxic injection details) into V1 of SST-Cre mice. After perfusion, sections were labeled with rabbit anti-somatostatin antibodies (1:1000; Peninsula Laboratories) and the SST interneuron-specific GFP expression was confirmed. V1 was micro-dissected from an additional cohort of mice to assay overexpression of Lypd6 through qPCR. RNA was isolated using a RNeasy lipid tissue mini kit (Qiagen) and cDNA was produced. The cDNA was subjected to qPCR analysis using a Taqman assay (Life Technologies) at the Icahn School of Medicine at Mount Sinai Quantitative PCR CORE facility.

### Stereotaxic injection

Mice were isoflurane anesthetized and head-fixed in a mouse stereotaxic apparatus (Narishige). The age range for surgery was P60 minimum to P78 maximum with median age at P64. A mid-line incision was made in the scalp and a micro-drill was used to drill a small hole in the skull over the right visual cortex. Three injections (0.5 μl each) were made into the deep layers of V1 binocular zone (from lambda: AP: 0.0, ML: 3.1, DV: 0.6; AP: 0.0, ML: 2.85, DV: 0.6; AP: 0.3, ML: 3.0, DV: 0.6) through glass micropipettes (1.14mm diameter, 20∼30um at opening) mounted to a Nanoject III Nanoliter Injector (Drummond Scientific) at 1nl/second. The syringe remained in place for five minute following injection to reduce backflow of virus. After injections, the skull hole was sealed using Kwik-Sil (World Precision Instruments), and the scalp was sutured. The mice recovered from anesthesia in an empty cage over a warming pad. Following recovery, mice were returned to their home cage where they remained for >3 weeks to allow for viral incorporation prior to any additional procedures or testing. The following viral constructs were used in this study: AAV-Lypd6 (AAV8-EF1α-DIO-EGFP-2A-Lypd6), AAV-GFP (AAV8-hSyn-DIO-EGFP: University of North Carolina (UNC) Vector Core), AAV-mCherry (AAV8-hSyn-DIO-mCherry: Addgene), AAV-CamKII-Cre (AAV8-CamKIIa-mCherry-Cre: UNC Vector Core), AAV-GqDREADD (AAV8-hSyn-DIO-hM3D(Gq)-mCherry: UNC Vector Core), AAV-ChR2 (AAV1-Ef1α-dflox-hChR2(H134R)-mCherry-WPRE-hGH: Penn Vector Core/Addgene), and AAV-hSyn-Cre (AAV8-hSyn-mCherry-Cre: UNC Vector Core).

### Chemogenetic activation of hM3d(Gq) Designer Receptors Exclusively Activated by Designer Drugs (DREADD)

Clozapine-N-Oxide (CNO; Sigma-Aldrich), a normally inert compound that specifically activates DREADD receptors, was prepared in 0.9% saline and injected I.P. into adult SST-Cre **(Fig. 4E-H)** at a concentration of 3 mg/kg. CNO or saline was injected immediately following MD and again 12 hours later for DREADD induced activation of SST interneurons during the first day of MD following the previously validated protocol in mouse visual cortex(Kuhlman et al., 2013).

**Fig. 1.**
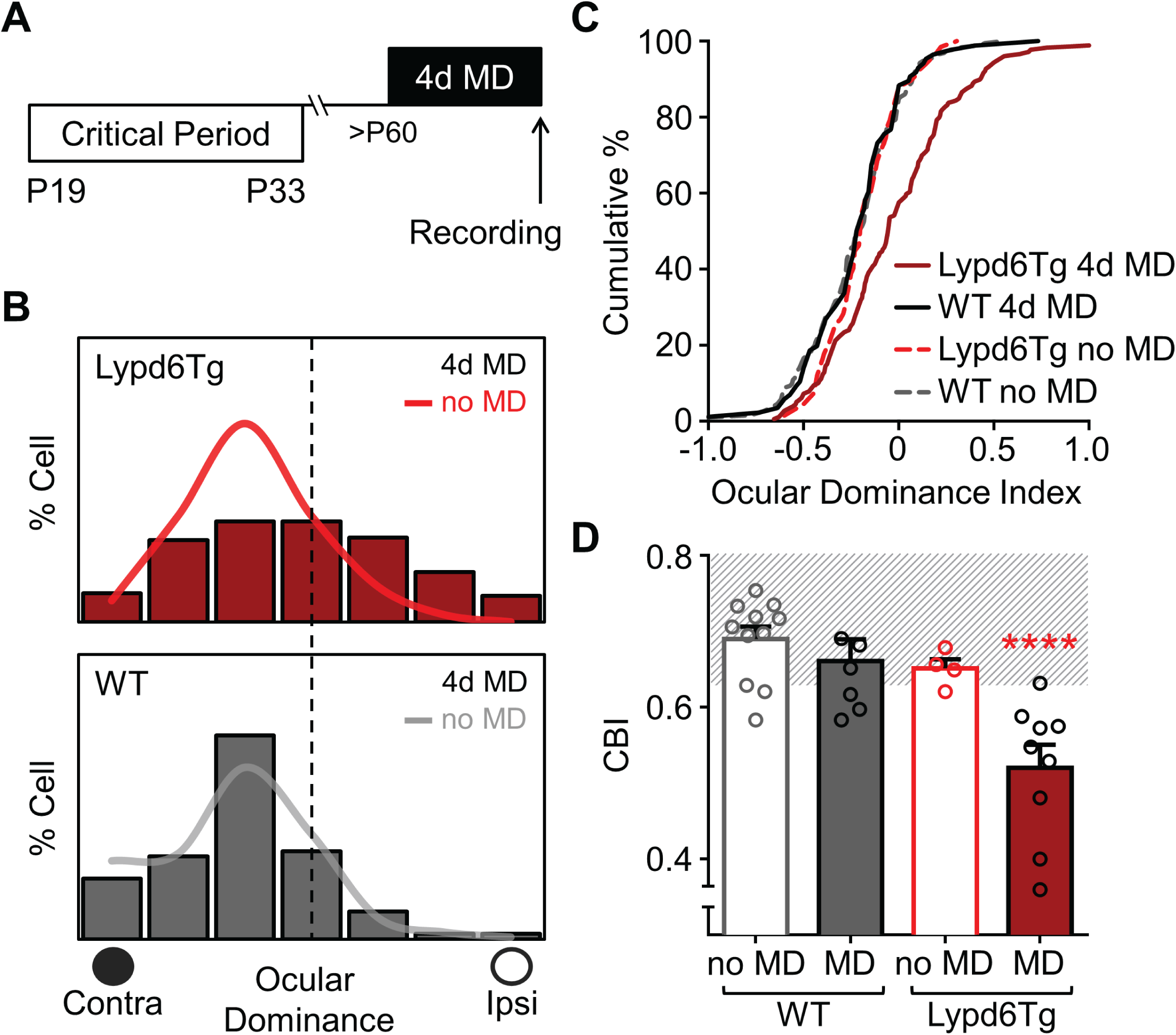
Neuronal overexpression of Lypd6 prolongs ocular dominance plasticity into adulthood. **(A)** Schematic of V1 plasticity paradigm with 4 days of monocular deprivation (4d MD) ending with in vivo electrophysiological recording from the V1 binocular zone in adult (>P60) Lypd6Tg or WT mice. **(B)** Adult 4d MD results in a shift in ocular dominance distribution of Lypd6Tg mice [red bar histogram; n=179 cells from 9 mice] but not of WT mice [gray bar histogram; n=86 cells from 7 mice], Lypd6Tg no MD [bright red line histogram; n=67 cells from 5 mice] and WT no MD [gray line histogram; n=193 cells from 11 mice]. Statistics based on actual cell numbers recorded, but histograms represent percentage of cells with each ocular dominance scores. Main groups are presented in bar histogram format. Control groups are overlaid as line format. Filled circle labeled as “contra” represents contralateral eye that received monocular deprivation in 4d MD designated groups. Empty circle labeled as “ipsi” represents the ipsilateral non-deprived eye. **(C)** Cumulative plot of ocular dominance index (ODI) after adult 4d MD confirms ocular dominance shift in Lypd6Tg mice [red line; n=179 cells from 9 mice] but not in WT mice [black line; n=86 cells from 7 mice], Lypd6Tg no MD [dashed bright red line; n=67 cells from 5 mice], and WT no MD [dashed gray line; n=193 cells from 11 mice]. **(D)** Comparison of contralateral bias index (CBI) following adult 4d MD in WT mice [gray solid bar; CBI=0.66, n=7 mice] and Lypd6Tg mice [red solid bar; CBI=0.52, n=9 mice], or no MD in WT mice [gray open bar; CBI=0.66, n=11 mice] and Lypd6Tg mice [red open bar; CBI=0.65, n=5 mice]. Lypd6Tg 4d MD has significantly decreased CBI compared to all other groups: WT no MD, WT 4d MD, and Lypd6Tg no MD. Striped background area represents CBI range of an adult WT mouse. Data in figure presented as mean±SEM.

**Fig. 2.**
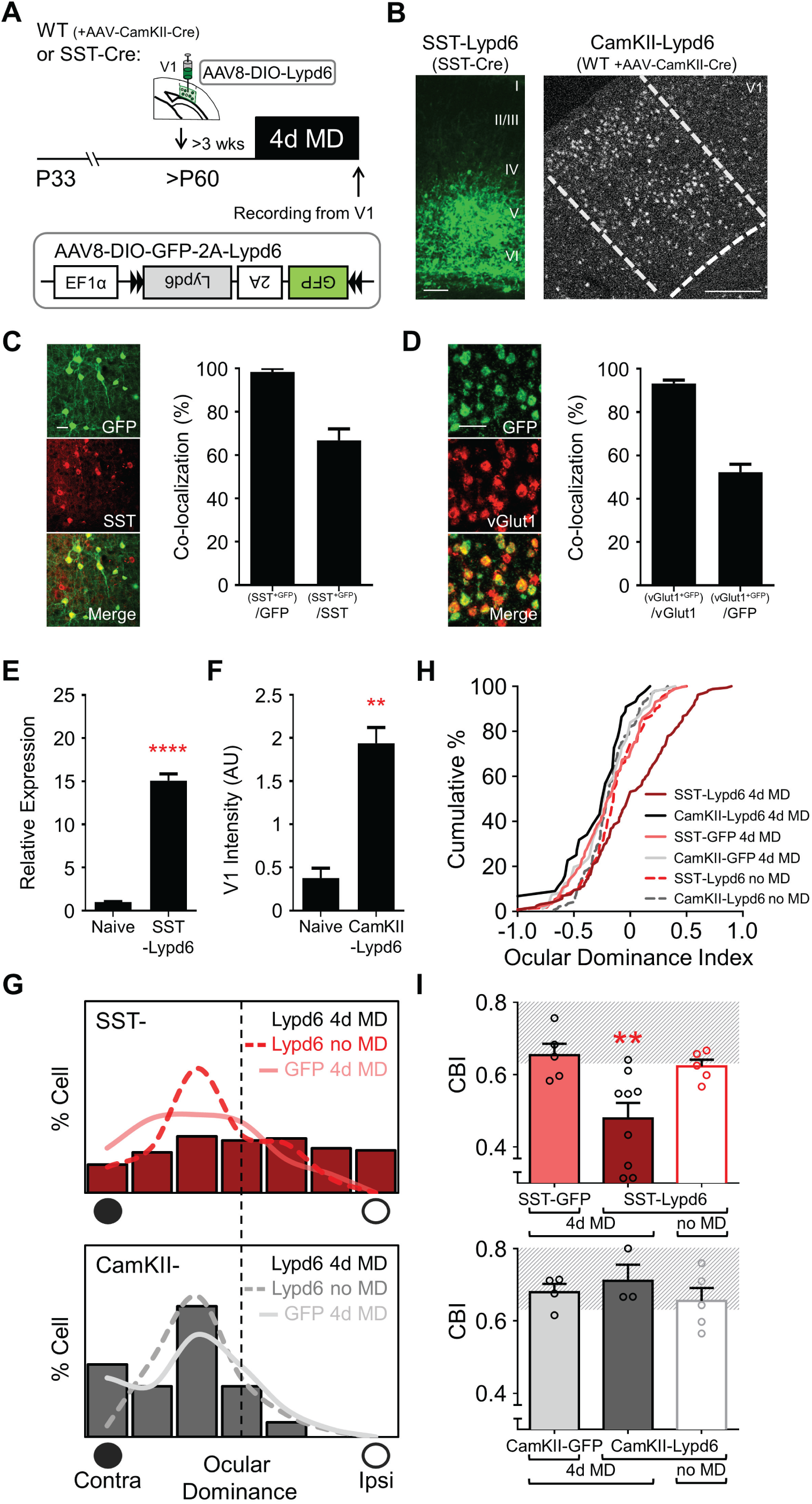
Selective overexpression of Lypd6 in SST interneurons reactivates ocular dominance plasticity in adult V1. **(A)** Schematic of 4d MD V1 plasticity paradigm with V1 stereotaxic injection of viral Lypd6 overexpression construct. AAV8-DIO-GFP-2A-Lypd6 (see inset) was injected into the binocular zone of V1 of adult SST-cre mice, or WT mice (co-injected with AAV-CamKII-cre) and incubated for >3 weeks prior to 4d MD. **(B)** Left: representative image of V1 following injection of AAV-Lypd6 in SST-Cre mice (SST-Lypd6), where viral transduction is represented by GFP (green). Scale bar=100μm. Right: representative image of fluorescent *in situ* hybridization labeling of Lypd6 in V1 binocular zone following co-injection of AAV-Lypd6 and AAV-CamKII-Cre in WT mice (CamKII-Lypd6). Scale bar=200μm. **(C)** Representative immunohistochemistry in V1 binocular zone of SST-Lypd6 mouse showing viral GFP (green) overlapping immunolabeled SST (red) neurons. Scale bar=50μm. 98±1.7% of cells expressing viral GFP were immunolabeled with SST. 67±5% of SST immunolabeled cells co-expressed viral GFP. [n=2 mice, 4 sections each, total from 8 sections: 190 co-labeled cells, 194 GFP labeled cells, 280 SST labeled cells] **(D)** Representative double *in situ* hybridization labeling of viral GFP in V1 binocular zone of CamKII-Lypd6 mouse showing specific expression of GFP (green) in vGlut1 (red)-positive neurons. Scale bar=50μm. 93.2±1.6% of cells expressing GFP mRNA co-express vGlut1 mRNA. 52.3±3% of vGlut1 mRNA-positive cells co-express viral GFP mRNA. [n=3 mice, 2 sections each, total from 6 sections: 979 co-labeled cells, 1051 GFP labeled cells, 1879 vGlut1 labeled cells] **(E)** Quantitative PCR of *Lypd6* mRNA from cDNA derived from whole V1 extracts of naïve (non-injected) and AAV-Lypd6 injected SST-cre mice [ΔΔCT method, n=3 mice]. **(F)** Quantification of Lypd6 expression in V1 binocular zone via fluorescent *in situ*-labeled sections from CamKII-Lypd6 mice, compared between injected and naïve (non-injected) hemispheres. Absolute intensity for every Lypd6 positive cells in the binocular V1 was summed and the total value was subsequently divided by the μm^2^ area of images of V1 binocular zone. Higher Lypd6 expression in the CamKII-Lypd6 hemisphere [5 sections from 2 mice]. **(G)** 4d MD in SST-Lypd6 mice (SST-Lypd6 4d MD) results in ocular dominance shift [red bar histogram; n=158 cells from 9 mice] but not in CamKII-Lypd6 mice (CamKII-Lypd6 4d MD) [gray bar histogram; n=44 cells from 3 mice], SST-Lypd6 no MD [dashed bright red line histogram; n=110 cells from 5 mice], and SST-GFP 4d MD [pink line histogram; n=86 cells from 5 mice]. **(H)** Cumulative ODI plot confirms ocular dominance shift in SST-Lypd6 4d MD mice [red line; n=158 cells from 9 mice] but not in CamKII-Lypd6 mice [black line; n=44 cells from 3 mice], SST-Lypd6 no MD [dashed bright red line; n=110 cells from 5 mice], and SST-GFP 4d MD [pink line; n=86 cells from 5 mice]. **(I)** Comparison of CBI following 4d MD in SST-Lypd6 [red solid bar; CBI=0.48, n=9 mice], SST-GFP [pink solid bar; CBI=0.65, n=5 mice], CamKII-Lypd6 mice [dark gray solid bar; CBI=0.71, n=3 mice], and CamKII-GFP [light gray solid bar; CBI=0.68, n=4 mice], with non-deprived SST-Lypd6 [red open bar; CBI=0.62, n=5 mice] and non-deprived CamKII-Lypd6 [gray open bar; CBI=0.66, n=5 mice]. SST-Lypd6 4d MD significantly differs from: SST-Lypd6 no MD, SST-GFP 4d MD, and CamKII-Lypd6 4d MD. Data in figure presented as mean±SEM.

**Fig. 3.**
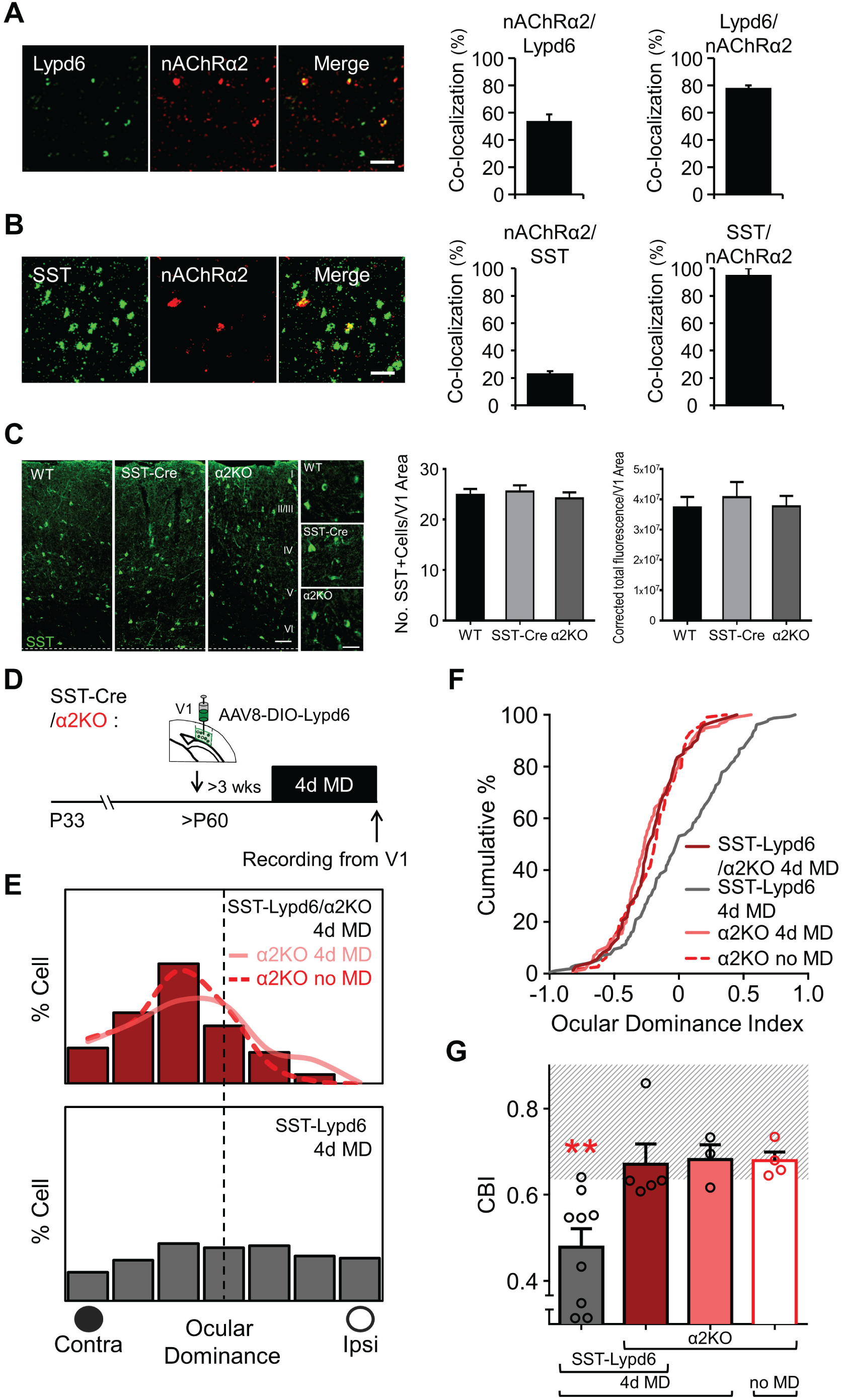
The α2 nicotinic acetylcholine receptor subunit is required for Lypd6-mediated reactivation of ocular dominance plasticity in adult V1. **(A)** Double fluorescent labeling of mRNA in layer V of adult V1 for *Lypd6* (green) and *α2 nicotinic acetylcholine receptor (Chrna2: nAChRα2)* (red). 53±3.26% of *Lypd6* mRNA labeled cells co-express *nAChRα2*. 79±2.93% of *nAChRα2* mRNA labeled cell co-express *Lypd6* mRNA. [n=2 mice, 8 sections]. **(B)** Double fluorescent *in situ* hybridization labeling of mRNA in layer V of V1 for *SST* (green) and *nAChRα2* (red). 23±0.01% of *SST* mRNA labeled cell co-express *nAChRα2* mRNA. 95±0.04% of *nAChRα2* mRNA labeled cells co-express *SST* mRNA. [n=2 mice, 4 sections per mouse]. Scale bars=50μm. **(C)** Comparison of immunohistochemical labeling (green) of somatostatin (SST) in V1 binocular zone of wild type (WT), SST-Cre and Chrna2KO (α2KO) mice. Scale bar=100μm (macro), 50μm (close-up). (Left graph) Number of SST+ cells per V1 area. (Right graph) Corrected total fluorescence per V1 area. WT, SST-Cre and Chrna2KO mice have comparable total number of V1 somatostatin-positive cells or levels of somatostatin immunopositivity [n=3 mice per group, 3 sections per mouse, group total: WT=112 cells, SST-Cre=115 cells, α2KO=109 cells]. **(D)** AAV-Lypd6 was injected into the V1 binocular zone of adult bigenic SST-cre/Chrna2KO mice (SST-Lypd6/α2KO), or adult SST-cre mice (SST-Lypd6) and incubated for >3 weeks prior to 4d MD. **(E)** Ocular dominance shift is reduced in bigenic SST-Lypd6/α2KO 4d MD mice [red bar histogram; n=73 cells from 5 mice] compared to SST-Lypd6 4d MD mice [gray bar histogram; data in Fig. 2G legend]. 4d MD does not affect ocular dominance in adult Chrna2KO mice [pink line histogram; n=116 from 3 mice] (α2KO no MD [red dotted line histogram; n=114 from 4 mice]). **(F)** Cumulative ODI plot confirms significantly reduced ocular dominance shift in the absence of nAChRα2 (SST-Lypd6/α2KO 4dMD) [red line; n=73 cells from 5 mice] compared to SST-Lypd6 4d MD mice [gray line; data in Fig. 2H legend]. **(G)** Comparison of CBI following 4d MD in SST-Lypd6 mice [gray solid bar; data in Fig. 2I legend], SST-Lypd6/α2KO mice [red solid bar; CBI=0.66, n=5 mice], and α2KO mice [pink solid bar; CBI=0.68, n=3 mice], or no MD in α2KO mice [bright red open bar; CBI=0.68, n=4 mice]. SST-Lypd6/α2KO 4d MD is only significantly different with SST-Lypd6 4d MD. Data in figure presented as mean±SEM.

**Fig. 4.**
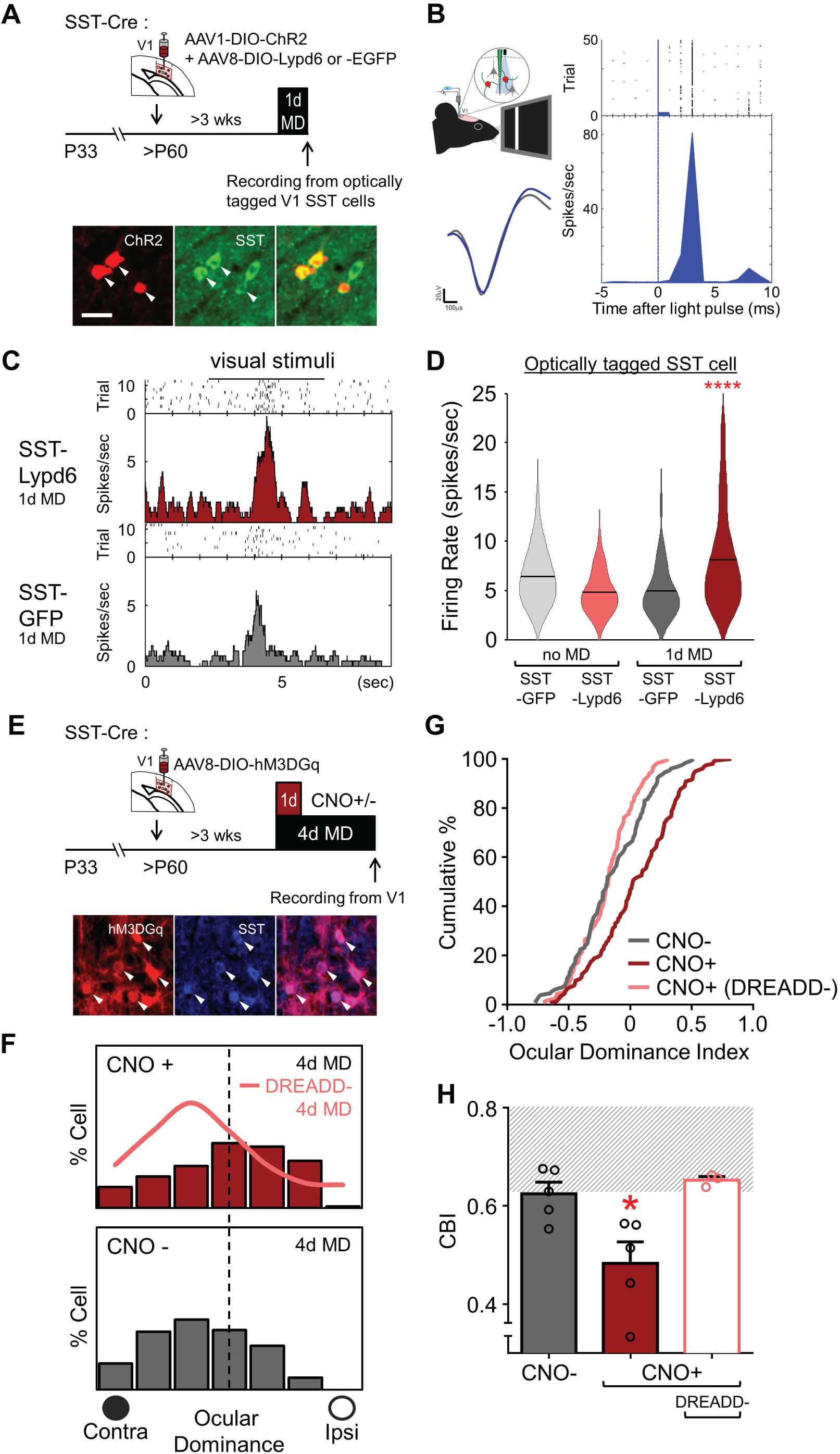
Lypd6 overexpression rapidly increases SST interneuron activity to express ocular dominance plasticity. **(A)** Adult SST-cre mice were injected with a combination of cre-dependent AAV-ChR2 and either AAV-Lypd6 (SST-Lypd6) or AAV-EGFP (SST-GFP) into the V1 binocular zone. Visually evoked firing rates of optically-tagged SST interneurons were recorded after 1 day of monocular deprivation (1d MD). Bottom: Representative image showing viral ChR2/mCherry (red) overlapping immunolabeled SST (green) neurons. 95±1.86% of ChR2/mCherry labeled cells co-labeled with SST [n=2 mice, 177 of 187 cells; mean±SEM]. Scale bar=25μm. **(B)** Top left: Schematic of *in vivo* electrophysiology setup with optical tagging. Bottom left: Representative traces of a putative SST interneuron from optical stimulation (blue trace) and averaged visually evoked spike (gray trace). Right: Histogram and raster plot shows time locked optogenetic activation of optically-tagged SST interneuron by 473 nm light pulses (20Hz, 1ms) delivered through optic fiber positioned above the recording site and coupled to the multichannel linear electrode. **(C)** Representative histograms of visually evoked firing of optically-tagged SST interneurons after 1d MD in SST-Lypd6 [red, top] and SST-GFP [gray, bottom]. **(D)** Comparison of visually evoked firing rate of SST interneurons in deep layer V1 following adult 1d MD in SST-GFP [gray solid bar; n=68 cells from 4 mice] and SST-Lypd6 mice [red solid bar; n=101 cells from 5 mice] with non-deprived (no MD) SST-GFP [gray open bar; n=82 cells from 5 mice] and SST-Lypd6 mice [red open bar; n=38 cells from 3 mice]. 1d MD results in significant increase in visually evoked firing rate under Lypd6 overexpression (SST-Lypd6 1d MD) in comparison to controls: SST-GFP 1d MD, SST-GFP no MD, and SST-Lypd6 no MD. **(E)** AAV-GqDREADD was injected into V1 binocular zone of adult SST-cre mice and incubated for >3 weeks prior to 4d MD. CNO (CNO+) or saline (CNO-) was given to only first day of 4d MD. **(F)** Chemogenetic activation of SST interneurons during the first day of 4d MD [CNO+: red, n=154 cells from 5 mice] results in significant ocular dominance shift, but not in mice given saline [CNO-: gray, n=129 cells from 5 mice], or 4d MD mice that received CNO without expressing DREADD [DREADD-:pink line histogram; n=117 cells from 3 mice]. **(G)** Cumulative ODI plot confirms ocular dominance shift after 4d MD in +CNO [red line, n=154 cells from 5 mice] compared with CNO- [gray line, n=129 cells from 5 mice], or DREADD- [pink line; n=117 cells from 3 mice]. **(H)** Comparison of CBI following 4d MD in CNO+ [red solid bar; CBI=0.50, n=5 mice], CNO- [gray solid bar; CBI=0.62, n=5 mice], and DREADD- [open pink bar; CBI=0.65, n=3 mice]. CNO+ significantly differs from: CNO- and DREADD-. Data in figure presented as mean±SEM.

### *In vivo* extracellular electrophysiology

Preparatory surgery leading to, and recording itself was performed initially under Nembutal/chlorprothixene anesthesia and then maintained with isoflurane (Bukhari et al., 2015). For animals that underwent either 4 days or 1 day of monocular deprivation, the deprived (contralateral) eye was reopened just prior to the start of recordings. Visually evoked single-unit activity was recorded in response to a high contrast single bar, generated by ViSaGe Stimulus Generator (Cambridge Research Systems), that laterally traveled across the monitor in a 4 second interval followed by a 6 second interstimulus interval per trial. Using an eye patch, the stimulus was separately presented to the contralateral eye and then to the ipsilateral eye for at least 12 trials each. For each animal, 3 to 10 single units were recorded in each of the 4 to 6 vertical penetrations spaced evenly (250 μm intervals) across mediolateral extent of V1 to map the monocular and binocular zones to avoid sampling bias. A linear 16-channel electrode was used to record each vertical penetration, which allowed recording from upper (channels 1∼8) and lower (channels 9∼16) cortical layers at similar weights. The signal detected from the probe was amplified and thresholded, and unit-sorted using an OmniPlex Neural Recording Data Acquisition System (Plexon). By comparing experimental conditions from 7 of the experimental groups that were used in our data, we found that the fraction of neurons recorded from upper layers (0.5235 ± 0.02350) did not differ from that of neurons recorded from lower layers (0.4765 ± 0.02350) (n=35 animals: *P*=0.3247, Student’s t-test for paired samples). A line crossing or “geometric” online sorting method was used to capture visually evoked responses as units. To ensure single-unit isolation, the waveforms of recorded units were further examined using Offline Sorter (Plexon). To analyze the electrophysiology data, normalized OD index of single neuron was computed by custom made MATLAB (MathWorks) script. This program generates peristimulus time histograms from each trial of visual stimulus presentation and uses a sliding window method to calculate maximum response during stimulus presentation, and then baseline (spontaneous) spiking activity by bootstrapping with the same sized sliding window during pre-stimulus. The program then compares the mean maximum visually evoked response to the mean baseline spiking activity using paired t-test to verify the significance of the visually evoked response. The analyzed peak to baseline spiking activity in response to each eye was then used to calculate OD scores using the formula: [[Peak(ipsi)-baseline (ipsi)]–[Peak (contra)-baseline(contra)]]/[[Peak (ipsi) – base line(ipsi)]+ [Peak(contra)-baseline(contra)]].). OD scores were converted from OD index using a 7-point classification scheme as follows: −1 to −0.5 = 1, −0.5 to −0.3 = 2, −0.3 to −0.1 = 3, −0.1 to 0.1 = 4, 0.1 to 0.3 = 5, 0.3 to 0.5 = 6, 0.5 to 1 = 7. For each binocular zone, contralateral bias index (CBI) is calculated according to the formula: [(n1-n7)+2/3(n2-n6)+1/3(n3-n5)+N]/ 2N, where N=total number of cells and nx=# of cells corresponding to OD score of x.

In experiments where visually evoked firing rates of SST-or PV interneurons were analyzed, a PCA cluster-based spike shape template-based sorting method was used. Visually evoked spikes assumed to be spikes originating from the same neuron grouped into visually identifiable subsets of waveforms to form clusters on 2D PCA space. Spike shape templates were selected by manually drawing selection contours around PCA clusters. The selected spikes were then automatically averaged and used to create templates, which were then used to sort incoming visually evoked spikes of similar shape and placement in PCA space into units. This online method allows defining multiple visually responsive units on a single channel of the linear 16-channel electrode, allowing recording up to 175 total cells in the V1 across 4∼6 sampling sessions. The number of putative PV interneurons or narrow-spiking (NS) cells recorded per sampling across all animals from all groups ranged between 11- 46 cells. Total cells (all cell types included) recorded, the number ranged from 53-172. In agreement with previously reported sorting methods (Niell and Stryker, 2008; Stark et al., 2013), the fractions of NS cells recorded were between 17.4∼24.2% of the total neurons recorded (SST-GFP 1d MD: 24.2%, SST-Lypd6 1d MD: 18.9%, SST-GFP no MD: 17.4%, SST-Lypd6 no MD: 20.5%).

For the experiment where SST interneurons were optogenetically tagged, a Cre-dependent virus expressing channelrhodopsin-2 (AAV1-Ef1α-dflox-hChR2(H134R)-mCherry-WPRE-hGH: Penn Vector Core) was injected into the V1 binocular zone of SST-Cre mice. The expression of channelrhodopsin in a cell type of interest allowed the use of optogenetic stimulation with a blue light as a search stimulus to simultaneously sort stimulus-locked responses from cell types of interest, and then record their visually evoked responses, by using an optic fiber-coupled 16-channel linear silicone probe (Neuronexus). This allowed accurate and high-throughput recordings of specific cell types, even against broad baseline noise and activities of other neuronal populations. Optogenetically tagged SST interneurons of the V1 binocular zone were identified using a 473 nm (blue) laser search stimulus emitted and delivered through the optic fiber coupled to the silicone probe and oriented immediately above the V1 cortical surface. After sorting for optogenetically responsive units (SST interneurons expressing ChR2), the optogenetic stimulus was switched off. Then a high-contrast single bar visual stimulus was presented to each eye to record the visually evoked responses (spike firing rate) of the sorted units. Analysis of visually evoked activities of sorted SST interneurons were restricted to only those recorded from the lower half of the 16-channel linear silicone probe to focus on the SST interneurons of lower layer IV, layers V, and VI. Recording from mice with less favorable conditions including suboptimal health and prepped brain conditions resulted in overall reduced visually evoked responses and subsequently reduced number of recorded neurons because of increased difficulty in recording. Because we deemed such animals would potentially misrepresent their groups, we established a measure of qualitative control by setting a quantitative threshold on number of recorded neurons per mouse to exclude recordings that were suboptimal. All animals with less than 10 analyzable lower layer SST interneurons recorded were excluded from the analysis. Normalized firing rate (baseline subtracted from peak firing rate) of each lower layer SST interneuron, in response to contralateral eye (or deprived eye in animals in 1 day monocular deprivation groups) stimulation, was computed by first using a custom made MATLAB script by generating a peristimulus time histogram-based analysis of peak and baseline spiking activity in response to visual stimulus. The peak visually evoked activity through contralateral/deprived eye stimulation was then subtracted by baseline spiking activity to obtain the normalized visually evoked firing rate.

To isolate visually evoked responses from narrow-spiking (NS) neurons, visual stimulus responsive cells were distinguished offline as NS cells or regular-spiking (RS) neurons by means of spike width- (trough-to-peak time) based classification. The spike-width criterion for separating into NS and RS neurons was established by measuring the spike width of optogenetically tagged PV interneurons. PV interneurons were optogenetically tagged by injecting a Cre-dependent virus expressing channelrhodopsin-2 (AAV1-Ef1α-dflox-hChR2(H134R)-mCherry-WPRE-hGH, Penn Vector Core) into V1 binocular zone of PV-Cre mice. During recording, NS cells were first identified and sorted, using a 473 nm laser search stimulus delivered through the optical fiber coupled to the 16-channel silicone probe and oriented immediately above the V1 cortical surface, depending on their responsiveness to blue light stimulation within 3 msec post blue light emission. The search stimulus was then exchanged to a visual stimulus. Visually evoked spike widths of the PV interneurons that were identified by optical tagging were then pooled to establish the official criterion for NS cells as having visually evoked spike width time (trough-to-peak time) of less than 412 μsec. The criterion was preliminarily tested in adult WT mice and NS neurons collected were confirmed as putative PV interneurons due to their similarly high visually evoked firing rate that was significantly higher in comparison to that of non-NS neurons. Averaged spike data for each unit was analyzed using a custom-made MATLAB script to obtain spike width time for each unit. All NS cells sorted and recorded were included in the analysis of their visually evoked activity. Under the same basis applied to animals recorded for visually evoked responses in SST interneurons, all animals with less than 10 analyzable NS cells recorded were excluded. Using the same method for analyzing SST interneurons (1 mouse from the SST-GFP 1d MD group and 2 mice from the SST-Lypd6 1d MD group), normalized firing rates of NS cells were calculated from responses to contralateral/deprived eye stimulation.

Following recordings, all AAV injected mice were transcardially perfused and the extent of GFP or mCherry signal was utilized to assess the viral transduction. Only mice that exhibited GFP or mCherry signal in the recorded V1 area were included for the analysis of ocular dominance plasticity or SST/NS cell activity. The experimenters performed while blind to experimental conditions such as viral constructs or CNO/saline administration, but not monocular deprivation.

### Visual evoked potential recording

Preparatory surgery leading to and recording itself was performed under Nembutal/chlorprothixene anesthesia and then maintained with isoflurane (Bukhari et al., 2015). A low-impedance tungsten electrode (FHC) was inserted into the V1 binocular zone near 3.00mm from lambda, and to a depth between 400∼500 μm from the pial surface where maximal visual evoked potential (VEP) amplitude can be measured in mice (Porciatti et al., 1999). Transient VEPs in response to horizontal sinusoidal gratings with abrupt contrast reversals (100%, 1Hz) of varying spatial frequencies between 0.05∼0.5 cycles/degree were measured from signals that were band-pass filtered (0.1∼100Hz) and amplified (AM Systems) and read using a custom software (National Instruments). VEP amplitudes from at least 30 events were averaged in synchrony with the stimulus contrast reversal. For each spatial frequency, between 4∼5 sessions of 30 events were sampled. Visual acuity was defined after VEP amplitudes were plotted against spatial frequency, and a non-linear regression (semi-log) curve was applied to extrapolate the zero-amplitude point. The age range for VEP recordings was P60 minimum to P74 maximum with median age at P68.

### *In situ* hybridization

The production of probes and methodology for *in situ* hybridization has been previously described(Demars and Morishita, 2014). Briefly, RNA probes including a fluorescein or digoxinogen (DIG) tag were generated and utilized to label *Lypd6, SST, vGlut1, GFP*, and *Chrna2* mRNA in 7 μm sections of V1 from fresh frozen brains of animals at P28 (CP) and >P60 (adult). To fluorescently label mRNA, anti-fluorescein/DIG-POD (1:2000; Roche) and anti-fluorescein/DIG-Alkaline phosphatase (AP) (1:1000; Roche) antibodies were used. POD-conjugated antibody labeling of mRNA was proceeded with TSA Plus DNP signal amplification (Perkin Elmer) and a subsequently labeling with anti-DNP-KLH-488 antibodies (1:1000; Life Technologies). AP-conjugated-antibody-labeled mRNA was stained using HNPP/Fast Red (Roche). Imaging was performed using an LSM780 confocal microscope (Zeiss). ImageJ was used to quantify the density of labeled pixels from each image or to examine co-localization using a color based thresholding method. For the quantification of Lypd6 *mRNA* expression across age in V1, pixel density (>2 standard deviations above mean intensity of full image field) was determined from low magnification images of V1 binocular zone using ImageJ software. For the quantification of co-localization of *Lypd6* or *SST* with *nAChRα2*, the number of cells positive for *Lypd6/SST* or *nAChRα2* in each image was determined using ImageJ by automated counting using a threshold of >2 standard deviations above background and limiting to particles of >40 μm cell diameter. To calculate the co-localization percentage, first color based thresholding was utilized in ImageJ to isolate and quantify the co-localized cells, then percentage was calculated by dividing the number of co-localized cells by the number of *Lypd6/SST* or *nAChRα2* positive cells in each image. For the quantification of co-localization of *GFP* and *vGlut1* mRNA, thresholding was first utilized to isolate and calculate number of particles representing *GFP* labeled cells (>40 μm cell diameter). Then the corresponding *vGlut1* labeled image was redirected to the mask retained from the analysis of *GFP* labeled image to calculate total number and then percentage of *GFP* labeled cells with co-localized *vGlut1* labeling. For the quantification of *Lypd6* mRNA expression in V1 binocular zone of mice injected with a cocktail of AAV-Lypd6 and AAV-CamKII-Cre, low magnification images of binocular V1 from the injected or the corresponding naïve hemisphere were analyzed for particles after thresholding. Absolute intensity for each particle was then measured as the product of mean intensity and area of particle. All absolute intensity values were summed to obtain total binocular V1 intensity, which was subsequently normalized by dividing by the μm^2^ area of the V1 binocular zone. The V1 binocular zone of each image was assessed using Paxinos and Franklin’s *The Mouse Brain in Stereotaxic Coordinates* (1997) as reference.

### Immunohistochemistry

Anesthetized mice were transcardially perfused with cold 4% paraformaldehyde (PFA) dissolved in 0.1M phosphate buffer. The brains were post-fixed in 4% PFA at 4 °C, and cryoprotected in 30% sucrose solution. The frozen brains were sectioned into 30-μm-thick coronal sections using a cryostat (CM3050, Leica). Free-floating sections were washed in tris-buffered saline (TBS), pH 7.5, and then blocked in 1% bovine serum albumin in TBST (0.25% Triton X-100 in TBS) for 1 h. The sections were incubated with mouse anti-parvalbumin (1:500; Swant), and rabbit anti-somatostatin (1:1000; Peninsula Laboratories). After primary antibody incubation the slices were washed in TBST, followed by secondary antibody incubation with Alexa fluor dyes (Thermo Fisher Scientific). Imaging was performed using a Zeiss LSM780 confocal microscope at 20 or 40X magnification. The V1 binocular zone of each image was assessed using Paxinos and Franklin’s *The Mouse Brain in Stereotaxic Coordinates* (1997) as reference. The investigator performing the analysis was blind to the animal genotype.

### Statistical analysis

The following statistical approaches were utilized for experiments assessing ocular dominance plasticity. χ^2^ test was used to compare at cell-level distributions of ocular dominance scores between two groups and assess for ocular dominance shift. For readability, the histogram figures for distribution of ocular dominance scores (OD Scores) represent percentage of cells rather than actual cell number. However, the χ^2^ statistics are results of the tests conducted on actual cell number. Cumulative distributions of ocular dominance index (ODI) were compared at cell-level by using the Kolmogorov-Smirnov (K-S) test. One-way analysis of variance (ANOVA) was used to compare contralateral bias index (CBI) at animal-levels and Tukey’s multiple comparisons test was used for post hoc analyses. As justification of the use of a parametric test, normality of the CBI data was tested by using the D’Agostino-Pearson Test and Shapiro-Wilk Test on any groups where n=mice exceeding the criteria for the tests (n>8, and n>7 respectively). Normality was confirmed in all groups where tests were applicable: WT MD, WT no MD, Lypd6Tg 4d MD, and SST-Lypd6 4d MD. In addition, we applied the Shapiro-Wilk Test to the CBI data of a key experimental group from our previous study (Smith et al., 2016) that utilized the same experimental design and statistical analyses for testing ocular dominance plasticity and confirmed normality. Comparisons of NS cell (PV interneuron) and SST interneuron firing rates were conducted using one-way ANOVA with Tukey’s multiple comparisons test for post hoc analyses. Student’s *t*-test was used for all other analyses that compare mean differences between two groups. A minimum *P* value of 0.05 was accepted as statistically significant. All statistical analyses were completed using Prism 6.0h (GraphPad Software). All CBI, quantified expression, quantified co-localization, and visually evoked firing rate data in figures are presented as mean±SEM. No method of randomization was used to determine allocation of samples. No statistical methods were used to predetermine sample sizes. However, all cell and animal-level sample sizes for all experiments including quantified expression/co-localization experiments and *in vivo* extracellular electrophysiological experiments were selected based on previously published studies.

## Results

### Neuronal overexpression of Lypd6 prolongs ocular dominance plasticity into adulthood

We began by testing whether Lypd6 can prolong heightened level of ocular dominance plasticity in adulthood beyond the V1 critical period. Lypd6 is known to potentiate calcium transmission through the nicotinic acetylcholine receptors (nAChR) (Darvas et al., 2009) in contrast to Lynx1, a fellow Ly6 family member known to limit nicotinic cholinergic signaling (Ibañez-Tallon et al., 2002; Miwa et al., 2006) and ocular dominance plasticity in adulthood (Morishita et al., 2010; Sadahiro et al., 2016). Therefore, we tested whether overexpression of Lypd6 would prolong ocular dominance plasticity into adulthood by using a transgenic mouse line that pan-neuronally overexpress Lypd6 under the human synapsin promotor (Lypd6Tg) (Darvas et al., 2009). We subjected adult Lypd6Tg mice to 4-days of monocular visual deprivation of the contralateral eye (MD), and then after the final day the deprived eye was reopened to assess ocular dominance in the V1 through *in vivo* extracellular electrophysiology under anesthesia (Gordon and Stryker, 1996; Bukhari et al., 2015) **(Fig. 1A)**. Each recorded V1 neuron was assigned an ocular dominance (OD) score based on the relative balance of the neuron’s visually evoked firing rates in response to visual stimuli independently presented to the contralateral and ipsilateral eye (Gordon and Stryker, 1996). Adult Lypd6Tg mice that underwent 4d MD displayed an overall significant shift in eye preference of the V1 (OD shift) away from the deprived contralateral eye and towards the non-deprived ipsilateral eye (Lypd6Tg 4d MD vs. WT 4d MD: χ^2^(6)=28.71, \*\*\*\**p*<0.0001). On the contrary, we did not detect OD shift in adult Lypd6Tg mice that were not subjected to monocular visual deprivation (no MD) (Lypd6Tg 4d MD vs. Lypd6Tg no MD: χ^2^(6)=20.87, \*\*\*\**p*<0.0001), or adult wild type (WT) mice that typically do not display plasticity regardless of 4 days of visual deprivation (Lypd6Tg no MD vs. WT no MD:]: χ^2^(6)=7.354, *p*=0.28, chi-squared test) **(Fig. 1B)**. Comparisons of cumulative distributions of ocular dominance index scores (ODI: −1=most ipsilateral dominant, 0=equal/binocular, 1=most contralateral dominant) for all recorded neurons from each group showed a significantly higher percentage of single neuronal responses that were ipsilateral dominant in the Lypd6Tg 4d MD mice (Lypd6Tg 4d MD vs. WT 4d MD: \*\*\*\**p*<0.0001; vs. Lypd6Tg no MD: \*\*\**p*=0.0002; Lypd6Tg no MD vs. WT no MD: *p*=0.2332, Kolmogorov-Smirnov (K-S) test) **(Fig. 1C)**. Lastly, individual animal level comparisons of contralateral bias index (CBI) scores – the relative strength to which the visually evoked activity of V1 neurons from the contralateral eye dominates over that from the ipsilateral eye, showed a significant decrease only in the Lypd6Tg 4d MD mice (F(3, 27)=10.70, \*\*\*\**p*<0.0001, one-way analysis of variance: ANOVA; Lypd6Tg 4d MD vs. WT no MD: \*\*\*\**p*<0.0001; vs. WT 4d MD: \*\**p*=0.0023; vs. Lypd6Tg no MD: \**p*=0.0208, Tukey’s multiple comparisons test) **(Fig 1D)**. Together, the results suggest overexpression of Lypd6 prolongs ocular dominance plasticity into adulthood and highlight a novel role of Lypd6 as a positive regulator of cortical plasticity.

### SST interneuron-selective Lypd6 overexpression reactivates ocular dominance plasticity in adulthood

While the prior experiment demonstrated the role of Lypd6 in prolonging ocular dominance plasticity into adulthood, the Lypd6 transgenic line cannot provide temporal, spatial and cell type details regarding this effect. In our previous study, we found that Lypd6 is enriched in V1 SST interneurons of the deep layers of the V1, and also to a lesser extent in a subset of glutamatergic neurons (Demars and Morishita, 2014). To characterize the selective effects of Lypd6 overexpression in adulthood in specific V1 neuronal subpopulations, we developed a Cre-dependent AAV with vector for overexpressing Lypd6 (AAV8-DIO-EGFP-2A-Lypd6: AAV-Lypd6). To achieve SST interneuron-specific overexpression, we injected AAV-Lypd6 into the binocular V1 of adult SST-Cre mice. Based on a previously implemented method (Koike et al., 2016; Pakan et al., 2016), we also overexpressed Lypd6 in V1 glutamatergic neurons of adult WT mice through viral cocktail injections of AAV-Lypd6 and a Cre expression AAV vector driven by a CamKII promoter (AAV8-CamKIIa-mCherry-Cre: AAV-CamKII-Cre) **(Fig. 2A)**. The viral injections in SST-Cre mice were adjusted to target the deep V1 areas containing layers V and VI where Lypd6 is endogenously expressed (Demars and Morishita, 2014) **(Fig. 2B)**. Both SST- and CamKII-targeted injection strategies resulted in robust target-specific viral expression. 98% of GFP labeled cells in adult SST-Cre mice co-labeled with SST while 67±5% of SST immunolabeled cells co-expressed viral GFP **(Fig. 2C)**. 93.2% of successfully co-transfected cells (GFP-positive as a result of both viral CamKII-selective Cre and Cre-dependent Lypd6 expression) in adult WT mice were positive for the excitatory cell marker vGlut1 while 52.3±3% of vGlut1-positive cells co-expressed viral GFP **(Fig. 2D)**. Overall significantly increased Lypd6 mRNA was measured across the V1 in both SST-selective (t(4)=17.67, \*\*\*\**p*<0.0001, Student’s *t*-test) and CamKII-selective Lypd6 overexpression (t(4)=3.742, *\*\*p*=0.0049, paired Student’s *t-*test) **(Fig. 2E, F)**.

We assessed whether adult overexpression of Lypd6 in SST interneurons influence ocular dominance plasticity. We injected AAV-Lypd6 or control AAV vector (AAV8-hSyn-DIO-EGFP: AAV-GFP) into the V1 of adult SST-Cre mice and, after >3 weeks of viral incorporation, subjected them to 4d MD. In adult mice with SST-specific Lypd6 overexpression, 4d MD (SST-Lypd6 4d MD) resulted in a significant OD shift (SST-Lypd6 4d MD vs. CamKII-Lypd6 4d MD: χ^2^(6)=29.72, \*\*\*\**p*<0.0001). On the contrary, we did not detect OD shifts in non-deprived counterparts (vs. SST-Lypd6 no MD: χ^2^(6)=31.79, \*\*\*\**p*<0.0001) or control 4d MD mice that were injected with AAV-GFP (vs. SST-GFP 4d MD: χ^2^(6)=23.77, \*\*\**p*=0.0006, chi-squared test) **(Fig. 2G)**. The cumulative distributions of ODI showed a significantly higher percentage of responses that were ipsilateral dominant only in the SST-Lypd6 4d MD mice (SST-Lypd6 4d MD vs. CamKII-Lypd6 4d MD: \*\*\*\**p*<0.0001; vs. SST-Lypd6 no MD: \*\*\**p*=0.0001; vs. SST-GFP: \*\*\**p*=0.0002, K-S test) **(Fig. 2H)**. Lastly, CBI was significantly decreased only in the SST-Lypd6 4d MD mice (F(5, 25)=5.732, \*\**p*=0.0012, one-way ANOVA; SST-Lypd6 4d MD vs. SST-Lypd6 no MD: \*\**p*=0.0477; vs. SST-GFP 4d MD: \*\**p*=0.0183, vs. CamKII-Lypd6 4d MD: \*\*\**p*=0.0073, Tukey’s multiple comparisons test) **(Fig. 2I)**. Because Lypd6 is also endogenously expressed in a subset of glutamatergic neurons, we also tested whether selective overexpression of Lypd6 in glutamatergic cells would induce plasticity in adult mice by injecting a viral cocktail of AAV-CamKII-Cre and either AAV-Lypd6 or AAV-GFP into V1 of adult WT mice. We did not observe a significant OD shift in mice with glutamatergic overexpression of Lypd6 that underwent 4d MD (CamKII-Lypd6 4d MD) **(Fig. 2G)**. Neither the distributions of OD scores nor CBI were different from those of control counterparts (CamKII-Lypd6 no MD and CamKII-GFP 4dMD) **(Fig. 2H, I)**. Altogether, robust ocular dominance plasticity was measured only in the visually deprived adult mice that overexpressed Lypd6 specifically in SST interneurons. These results suggest Lypd6-mediates reactivation of experience-dependent plasticity in adulthood through SST interneurons.

### Genetic deletion of α2 nicotinic acetylcholine receptor subunit abolishes Lypd6-mediated reactivation of ocular dominance plasticity

We next aimed to determine the nAChR subtype required for reactivation of V1 plasticity mediated by Lypd6 in SST interneurons. Aside from the deep cortical layer subpopulation of SST interneurons (Demars and Morishita, 2014), Lypd6 is notably expressed in a subpopulation of hippocampal SST-positive oriens-lacunosum moleculare (O-LM) interneurons, which are physiologically similar to cortical SST-expressing Martinotti-type interneurons (Heys et al., 2012). O-LM interneurons are also distinguished by the expression of the α2-subunit-containing nAChR (nAChRα2) – the subtype that is most sparsely expressed across the brain and features unique non-desensitizing receptor kinetics (Jia et al., 2010; Demars and Morishita, 2014). In the cortex, the expression of nAChRα2 is near-exclusively limited to the layer V Martinotti-type SST interneurons (Ishii et al., 2005; Tasic et al., 2016; Hilscher et al., 2017; Tasic et al., 2018). *In situ* hybridization for *Lypd6* and *Chrna2* mRNA revealed that 79% of nAChRα2-positive cells co-express Lypd6, and more than 53% of Lypd6-positive cells co-express nAChRα2 **(Fig. 3A)**. Furthermore, while 23% of SST-positive cells co-express nAChRα2, 95% of nAChRα2-positive cells co-express SST **(Fig. 3B)**. Consistent with recent single cell transcriptomic studies in adult V1 (Tasic et al., 2016; Tasic et al., 2018), no overlaps were observed between nAChRα2 and VIP or vGlut1 – the markers of vasoactive intestinal peptide-expressing interneurons and pyramidal cells (data not shown). Together, this confirms strong preferential co-expression of Lypd6 and nAChRα2 in a deep cortical layer subpopulation of SST interneurons.

We investigated whether experience-dependent plasticity after SST interneuron-specific overexpression of Lypd6 in adult mice requires nAChRα2. To accomplish this, we created a bigenic SST-Cre/Chrna2KO mouse line to allow SST-specific Cre-recombinase expression on a background of *Chrna2* knockout (Chrna2KO: α2KO) (Lotfipour et al., 2013). Adult Chrna2KO mice have SST expression levels and number of SST-positive cells comparable to those of adult WT and SST-Cre mice (adult WT, SST-Cre, a2KO mice SST-positive cell numbers: F(1.119, 2.238)=0.6923, *p*=0.502; SST immunopositivity: F(2, 24)=0.2158, *p*=0.8075, one-way ANOVA) **(Fig. 3C)**. Therefore, if plasticity induced by SST-selective Lypd6 overexpression is absent in the bigenic mice, then it would suggest an association of Lypd6 and nAChRα2 in the induction of plasticity, and not a disruption of the SST interneuron population due to the genetic ablation of *Chrna2*. We injected AAV-Lypd6 into V1 of adult SST-Cre/α2KO bigenic mice, allowed >3 weeks for viral incorporation, and assessed ocular dominance plasticity after 4d MD **(Fig. 3D)**. Intriguingly, the loss of nAChRα2 (SST-Lypd6/α2KO 4d MD) eliminated the robust cortical plasticity induced through SST-specific Lypd6 overexpression in adult V1. Unlike the adult SST-Lypd6 4d MD mice, adult SST-Lypd6/α2KO 4d MD mice as well as deprived and non-deprived adult α2KO mice (α2KO 4d MD and α2KO no MD) did not show significant shifts in OD (SST-Lypd6 4d MD vs. SST-Lypd6/α2KO 4d MD: χ^2^(6)=28.44, \*\*\*\**p*<0.0001; vs. α2KO 4d MD: χ^2^(6)=41.49, \*\*\*\**p*<0.0001; vs. α2KO no MD: χ^2^(6)=48.51, \*\*\*\**p*<0.0001, chi-squared test) **(Fig. 3E)**, shifts in cumulative distributions of ODI (SST-Lypd6 4d MD vs. SST-Lypd6/α2KO 4d MD: \*\*\*\**p*<0.0001; vs. α2KO 4d MD: \*\*\*\**p*<0.0001; vs. α2KO no MD: \*\*\*\**p*<0.0001, K-S test) **(Fig. 3F)**, or decreases in CBI (F(3, 17)=6.069, \*\**p*=0.0053, one-way ANOVA; SST-Lypd6 4d MD vs. SST-Lypd6/α2KO 4d MD: \*\**p*=0.0209; vs. α2KO 4d MD: \*\**p*=0.0450, vs. α2KO no MD: \*\*\**p*=0.0259, Tukey’s multiple comparisons test) **(Fig. 3G)**. These findings suggest the requirement of nAChRα2 in the induction of cortical plasticity by Lypd6 in SST interneurons.

### Rapid experience-dependent increase in SST interneuron activity drives ocular dominance plasticity under Lypd6 overexpression

We next investigated the cell-autonomous impact of Lypd6 overexpression on SST interneuron activity that underlies reactivation of V1 plasticity. Engaging cholinergic signaling through Lypd6 overexpression may potentiate SST interneuron activity and in turn disinhibit excitatory neurons through inhibition of PV interneurons. This effect would match the hallmark event of V1 plasticity during juvenile critical period wherein visual deprivation immediately results in a transient reduction of PV interneuron activity and in turn a disinhibition of the excitatory network to mediate ocular dominance plasticity (Hengen et al., 2013; Kuhlman et al., 2013; Feese et al., 2018). Considering Lypd6 is a positive modulator of nicotinic signaling (Darvas et al., 2009), and SST interneurons are well-positioned to inhibit PV interneurons (Cottam et al., 2013; Hioki et al., 2013; Pfeffer et al., 2013), we hypothesized that rapid experience-dependent increase in the activity of SST interneurons drives OD plasticity achieved through adult SST-selective overexpression of Lypd6.

To evaluate how overexpression of Lypd6 impacts SST interneuron activity, we expressed channelrhodopsin-2 (ChR2) as an optogenetic tag to allow efficient identification of individual SST interneurons amongst a population of visually responsive neurons by using a blue light (473nm) as a search stimulus. Using this technique, visually evoked neurons sorted were confirmed as SST interneurons if they also responded to optogenetic stimulation. We targeted expression of ChR2 in SST interneurons by injecting an AAV with cre-dependent expression of ChR2 (AAV1-dflox-hChR2(h134R)-mCherry-WPRE-hGH: AAV-ChR2) into the V1 of adult SST–Cre mice along with either AAV-Lypd6 (SST-Lypd6) or AAV-GFP (SST-GFP) After >3 weeks of viral incorporation, the mice were subjected to just 1 day of monocular deprivation (1d MD) **(Fig. 4A, B)**. Considering the endogenous expression of Lypd6 is limited to deep cortical layers, we restricted our analyses to SST interneurons recorded from the lower half of the 16-channel linear silicone probe that correspond to layers IV-VI. There was a significant increase in the visually evoked firing rate of SST interneurons in adult SST-Lypd6 mice that underwent 1d MD (SST-Lypd6 1d MD), compared to 1d MD adult SST-GFP mice (SST-GFP 1d MD) or non-deprived adult SST-Lypd6 (SST-Lypd6 no MD) mice (F(3, 267)=13.45, \*\*\*\**p*<0.0001, one-way ANOVA; SST-Lypd6 1d MD vs. SST-GFP 1d MD, vs. SST-GFP no MD, and vs. SST-Lypd6 no MD: respectively, \*\*\*\**p*<0.0001, \**p*=0.02 and \*\*\*\**p*<0.0001, Tukey’s multiple comparisons test) **(Fig. 4C, D)**. When baseline (spontaneous) firing rates without visual stimulation were compared, only a trending increase was observed when comparing SST-Lypd6 1d MD mice (4.47 spikes/sec) and SST-GFP 1d MD mice (2.87 spikes/sec), and no significant experience-dependent effects were observed SST-Lypd6 no MD (4.77 spikes/sec) (data not shown). This result suggests Lypd6 overexpression leads to a rapid experience-dependent increase in visually evoked SST interneuron activity.

To determine whether rapid increase in SST interneuron activity alone could mediate enhanced experience-dependent plasticity in adulthood, we examined whether chemogenetic activation of SST interneurons in adult mice strictly during the first of the 4 days of monocular deprivation can reactivate OD plasticity **(Fig. 4E)**. Adult SST-Cre mice were injected with an AAV with Cre-dependent expression of excitatory Gq protein-Designer Receptors Exclusively Activated by Designer Drugs (AAV8-hSyn-DIO-hM3D(Gq)-mCherry: AAV-GqDREADD)(Urban and Roth, 2015). Immediately following the start of 4d MD, SST interneurons were activated for the first 24 hours through administration of Clozapine-N-Oxide (CNO). As a result, activation of SST interneurons in the adult V1 during the first of 4 days of monocular deprivation (CNO+) resulted in significant OD shift as opposed to control saline administered (CNO-) mice or mice given CNO but do not express DREADD receptors (DREADD-) (CNO+ vs. CNO-: χ^2^(12)=53.12, \*\*\*\**p*<0.0001; vs. DREADD-: χ^2^(6)=42.52, \*\*\*\**p*<0.0001, chi-squared test) **(Fig. 4F)**. The cumulative distributions of ODI show a significant shift in CNO+ mice and not in controls (CNO+ vs. CNO-: \*\*\*\**p*<0.0001; vs. DREADD-: \*\*\*\**p*<0.0001, K-S test) **(Fig. 4G)**. Comparisons of CBI scores show a significant decrease only when SST interneurons were activated (F(2, 10)=7.402, \**p*=0.0107, one-way ANOVA; CNO+ vs. CNO-, vs. DREADD-: respectively, \**p*=0.02, and \**p*=0.02, Tukey’s multiple comparisons test) **(Fig. 4H)**. Together, these results reveal a novel role for SST interneuron activity during the initial phase of visual deprivation in reactivating robust cortical plasticity in adulthood.

### SST interneuron-selective Lypd6 overexpression leads to rapid experience-dependent suppression of PV interneuron and disinhibition of pyramidal neurons

After confirming immediate experience-dependent activation of SST interneurons underlies OD plasticity mediated by Lypd6 overexpression, we investigated how this affects downstream local circuits. We measured visually evoked activity of PV interneurons and excitatory neurons after 1 day of monocular deprivation under SST-specific Lypd6 overexpression under the hypothesis that experience-dependent activation of SST interneurons through Lypd6 overexpression reduces PV interneuron activity and in turn disinhibits excitatory neurons. From adult SST-Cre mice injected with either AAV-Lypd6 (SST-Lypd6) or AAV-GFP (SST-GFP) and subjected to 1d MD **(Fig. 5A)**, visually evoked cells were sorted based on spike width (trough-to-peak time) into two categories. Putative PV neurons were defined by a narrow spike (NS) waveform that is electrophysiologically unique to PV interneurons. The criterion, or spike width cutoff of 412μs, to determine NS neurons was established by collecting visually evoked spike widths of optically tagged PV interneurons in PV-Cre mice injected with AAV-ChR2 **(Fig. 5B)**. The visually evoked firing rate of the NS neurons preliminarily sorted in WT mice via the criterion, were significantly higher than that of non-NS (non-putative PV) neurons and similar to that of optically-tagged PV interneurons (F(3, 123)=16.92 *\*\*\*\*p*<0.0001, one-way ANOVA; NS vs. non-NS: \*\*\**p=*0.0001, vs. non-PV: \*\*\**p=*0.0001, vs. PV: *p=*0.9954, Tukey’s multiple comparisons test) **(Fig. 5C)** (See Materials and Methods: *In vivo* extracellular electrophysiology). Additionally, pyramidal neurons were defined using a combination of spike width sorting and SST interneuron optical tagging to isolate non-tagged and non-NS (regular-spiking: RS) neurons.

**Fig. 5.**
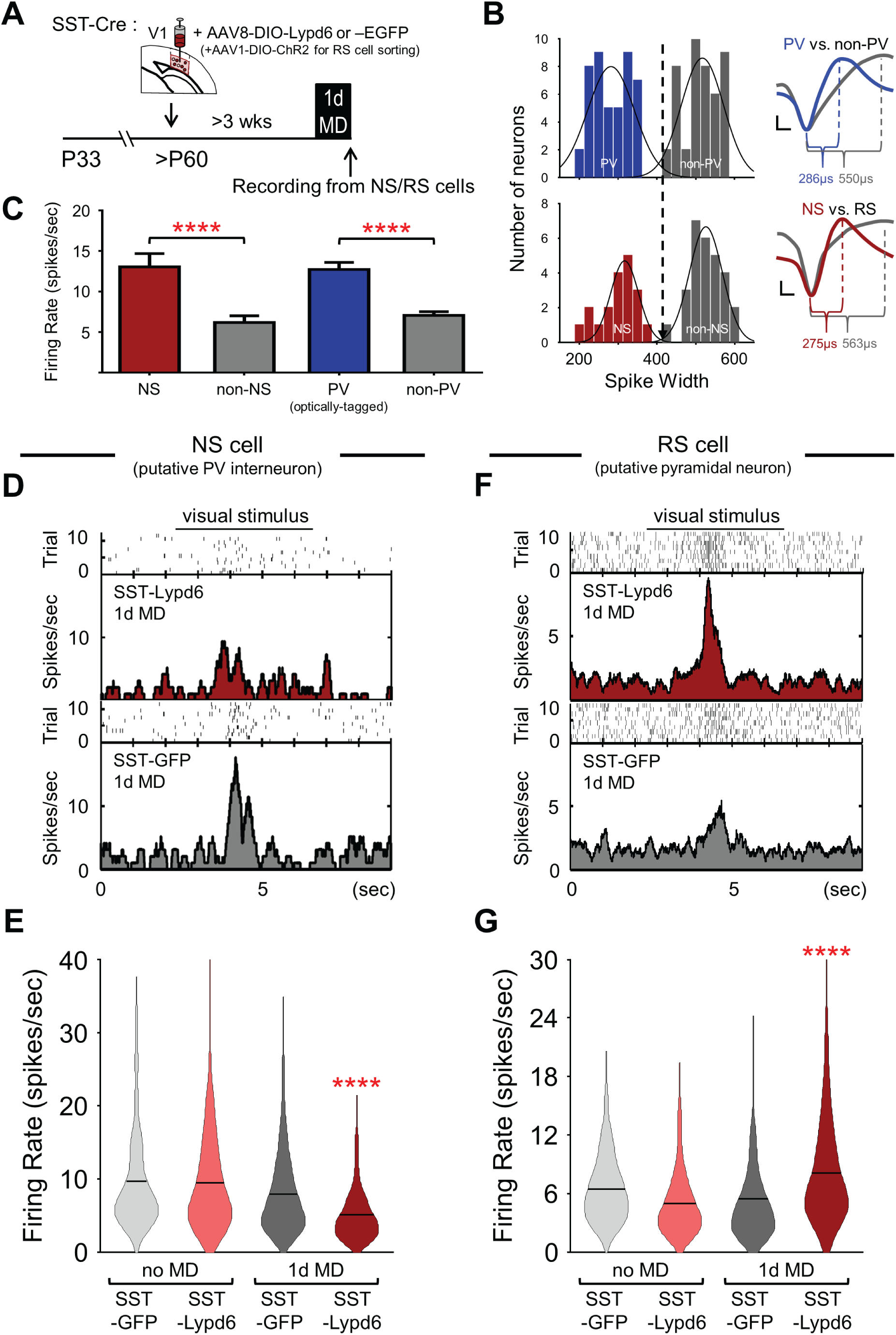
Rapid reduction in PV interneuron activity is required for ocular dominance plasticity under Lypd6 overexpression. **(A)** Adult SST-cre mice were injected with either AAV-Lypd6 (SST-Lypd6) or AAV-EGFP (SST-GFP) into the V1 binocular zone. For experiments involving RS cell (putative pyramidal neuron) sorting, AAV1-DIO-ChR2 was co-injected to exclude optically tagged SST interneurons. **(B)** Criterion for assessing narrow-spiking units as putative PV interneurons (NS cells) is defined as neurons with spike width (trough-to-peak time) less than 412μs (dotted line), established from visually evoked spike widths of optically tagged (ChR2-expressing) PV interneurons in PV-cre mice [top histogram, PV(Blue): 45 cells, non-PV(Gray): 45 cells, 3 mice]. The criterion was then preliminarily applied to visual evoked spikes collected from adult WT mice [bottom histogram, NS(Red): 19 cells, non-putative-PV interneuron (non-NS): 28 cells, 3 mice]. Right insets are representative traces of averaged visually evoked spikes from groups corresponding to the histograms (See details in Materials and Methods: *In vivo* extracellular electrophysiology). **(C)** Sorted NS cells have higher average visually evoked firing rate than non-NS and non-PV cells but are comparable to optically-tagged PV cells. **(D)** Representative histograms of visually evoked firing of NS cells after adult 1d MD in SST-Lypd6 [top, red] and SST-GFP [gray, bottom]. **(E)** Comparison of visually evoked firing rate of NS cells following adult 1d MD in SST-GFP [gray solid bar; n=148cells from 5 mice] and SST-Lypd6 [red solid bar; n=170 cells from 7 mice] with non-deprived (no MD) SST-GFP [gray open bar; n=116 cells from 5 mice] and SST-Lypd6 [red open bar; n=63 cells from 4 mice]. 1d MD results in significant decrease in visually evoked firing rate of NS cells under SST-selective Lypd6 overexpression (SST-Lypd6 1d MD) in comparison to controls: SST-GFP 1d MD, SST-GFP no MD, and SST-Lypd6 no MD. **(F)** Representative histograms of visually evoked firing of RS cells after adult 1d MD in SST-Lypd6 [top, red] and SST-GFP [gray, bottom]. **(G)** Comparison of visually evoked firing rate of RS cells following adult 1d MD in SST-GFP [gray solid bar; n=167 cells from 5 mice] and SST-Lypd6 [red solid bar; n=204 cells from 6 mice] with non-deprived (no MD) SST-GFP [gray open bar; n=240 cells from 6 mice] and SST-Lypd6 [red open bar; n=121 cells from 4 mice]. 1d MD results in significant increase in visually evoked firing rate under Lypd6 overexpression (SST-Lypd6 1d MD) in comparison to controls: SST-GFP 1d MD, SST-GFP no MD, and SST-Lypd6 no MD. Data in figure presented as mean±SEM.

In adult SST-Lypd6 mice that underwent 1 day of monocular deprivation of the contralateral eye (SST-Lypd6 1d MD), we found a significant reduction in visually evoked activity of NS neurons compared to all control groups, which suggests an experience-dependent decrease in firing rate of NS neurons when Lypd6 is overexpressed in adult SST interneurons (F(3, 488)=9.54, \*\*\*\**p*<0.0001, one-way ANOVA; SST-Lypd6 1d MD vs. SST-GFP 1d MD, vs. SST-GFP no MD, and vs. SST-Lypd6 no MD: all \*\*\*\**p*<0.0001, Tukey’s multiple comparisons test) **(Fig. 5D, E)**. Following that, we also found that only under SST-specific Lypd6 overexpression there is a significant experience-dependent increase in the visually evoked activity of non-SST RS neurons (F(3, 845)=23.60, \*\*\*\**p*<0.0001, one-way ANOVA; SST-Lypd6 1d MD vs. SST-GFP 1d MD, vs. SST-GFP no MD, and vs. SST-Lypd6 no MD: all \*\*\*\**p*<0.0001, Tukey’s multiple comparisons test) **(Fig. 5F, G)**. The contrasting experience-dependent changes in visually evoked activity of NS neurons and RS neurons amidst an increase in SST interneuron activity suggests that SST-specific Lypd6 overexpression may drive strong inhibition from SST to PV interneurons and in turn disinhibit the pyramidal neurons to rapidly fulfill the hallmark physiological event for initiating the cascade that results in reactivation of robust experience-dependent cortical plasticity in adulthood.

### SST interneuron-selective Lypd6 overexpression improves visual acuity after early long-term monocular deprivation

Finally, we tested whether enhanced cortical plasticity induced through SST-selective overexpression of Lypd6 can promote robust functional changes in adulthood and investigated this in the context of recovering lost visual acuity in the mouse model of amblyopia. When wild type mice are subjected to long-term monocular deprivation (LTMD) spanning the critical period, this results in persistent loss of visual acuity, which is measured directly in the V1 as the visual stimulus spatial frequency threshold that can evoke potential (VEP). Notably, visual acuity deficits endured after LTMD during the critical period cannot be recovered in adulthood even if the eye is reopened to restore and sustain visual input (Prusky and Douglas, 2003; Morishita et al., 2010). SST-Cre mice were subjected to LTMD between P19 through P33. Afterwards the deprived eye was reopened and maintained for the remainder of the experiment. At P45 the SST-Cre mice received V1 injections of either AAV-Lypd6 (SST-Lypd6) or control AAV-GFP (SST-GFP). Finally, after P60, the mice were tested for visual acuity of the previously deprived contralateral eye **(Fig. 6A).** Vial injections alone did not alter visual acuity, as all injected mice that were not subjected to early visual deprivation (no MD) had visual acuity that were comparable to that of adult SST-Cre mice naïve to both visual deprivation and viral conditions (naïve SST-Cre mice average visual acuity: 0.51 cycles/deg; data not shown in figure). Among the mice that underwent LTMD followed by an eye opening, only the adult SST-GFP mice displayed expected enduring deficits in visual acuity. On the other hand, the visual acuity of the adult SST-Lypd6 mice that underwent LTMD and the subsequent eye re-opening was significantly higher than that of the SST-GFP LTMD mice and was comparable to those of the non-deprived counterparts (F(3, 13)=29.91, \*\*\*\**p*<0.0001, one-way ANOVA; SST-GFP LTMD+eye reopened vs. SST-GFP no MD, SST-Lypd6 no MD, SST-Lypd6 LTMD+eye reopened: respectively, \*\*\*\**p*<0.0001, \*\*\**p*=0.0001, and \*\*\*\**p<*0.0001, Tukey’s multiple comparisons test) **(Fig. 6B, C)**. The normal level of visual acuity, despite LTMD, after adult overexpression of Lypd6 selectively in V1 SST interneurons suggests recovery of visual function, and suggests the capacity of plasticity induced through Lypd6 or SST interneurons to drive restoration of cortical function that is typically beyond reach in adulthood.

**Fig. 6.**
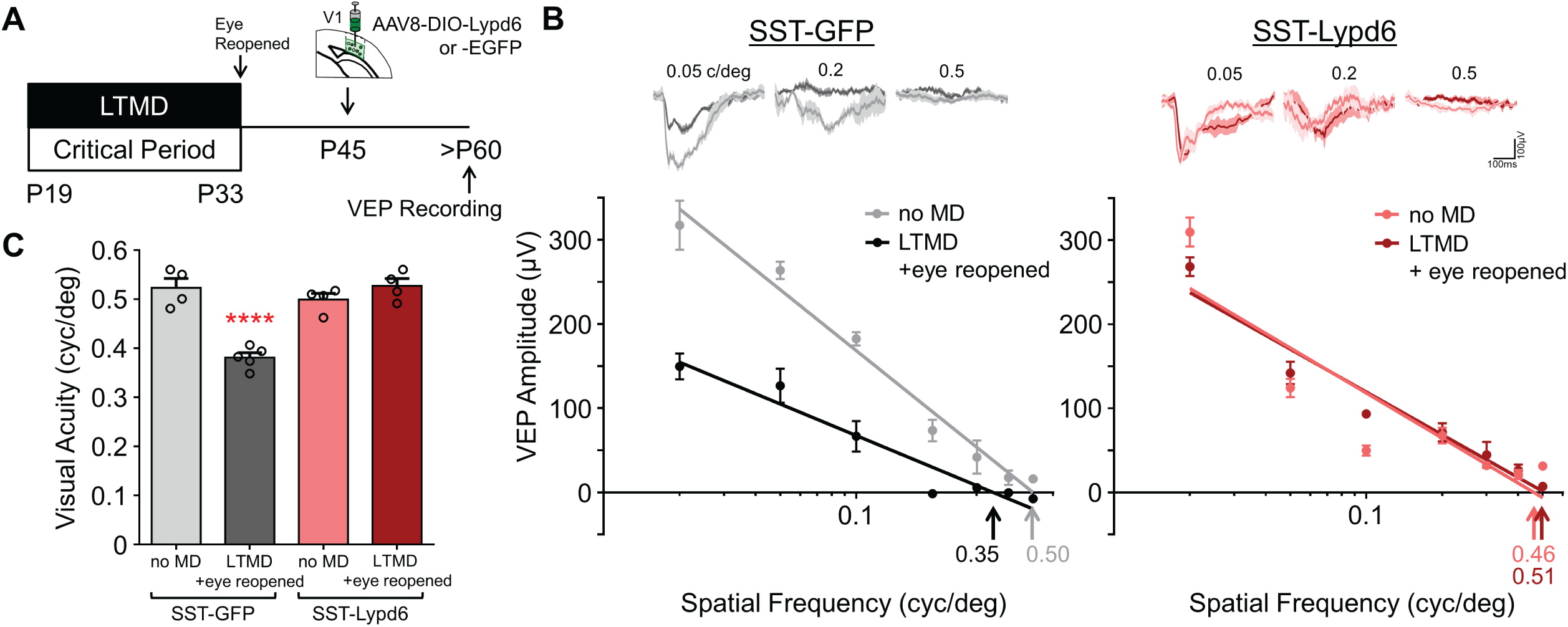
Functional recovery from amblyopia after V1 SST-selective overexpression of Lypd6. **(A)** SST-cre mice were subjected to long-term monocular deprivation (LTMD) from P19 until P33, then the deprived eye was reopened, and binocular vision was maintained for the remainder. At P45 AAV-Lypd6 (SST-Lypd6) was injected into the V1 binocular zone. Visual evoked potential (VEP) was assessed after P60. **(B)** Top: representative VEP traces in response to sinusoidal grating stimuli at 0.05 cyc/deg, 0.2 cyc/deg, 0.5 cyc/deg. Bottom: representative plots of VEP amplitudes of first negative peaks measured from a series of spatial frequencies in adult SST-Cre mouse overexpressing either Lypd6 (right series) or control GFP (left series) and with or without LTMD(+eye reopening): SST-GFP no MD (gray), SST-GFP LTMD+eye opened (black), SST-Lypd6 no MD (pink), and SST-Lypd6 LTMD+eye reopened (red). Arrows indicate visual acuity determined by extrapolated spatial frequency at 0V. **(C)** Comparison of visual acuity following LTMD+eye opening in SST-GFP mice [dark gray; Acuity=0.3809, n=5 mice] and SST-Lypd6 mice [red; Acuity=0.5315, n=4 mice], or no MD in SST-GFP mice [light gray; Acuity=0.5229, n=4 mice] and SST-Lypd6 mice [pink; Acuity=0.4990, n=4 mice]. SST-selective overexpression of Lypd6 post LTMD restores visual acuity to levels comparable to non-deprived mice. Only SST-GFP LTMD+eye reopened shows significantly perturbed visual acuity whereas SST-Lypd6 LTMD+eye reopened show visual acuity comparable to non-deprived controls. Data in figure presented as mean±SEM.

## Discussion

The decline of cortical plasticity after closure of the critical period is the key impedance of recovery from neurological disorders and brain trauma in later life. The current study identifies a molecule and circuit mechanism that modulate SST interneurons in the adult V1 to achieve rapid experience-dependent cortical disinhibition, one of the hallmarks of juvenile critical period plasticity, to enhance OD plasticity. We find that Lypd6, an endogenous nAChR modulator, enhances experience-dependent plasticity through the activation of adult cortical SST interneurons and disinhibition of excitatory neurons through suppression of PV interneurons. Intriguingly, increased cortical plasticity by overexpression of Lypd6 in SST interneurons promoted recovery from amblyopia during adulthood. This study also provides bases for future studies to investigate the mechanistic action and subcellular localization of Lypd6 in SST interneurons. Furthermore, our findings may provide insight for development of therapeutic compounds for treating brain disorders.

Like several of its fellow members of the Lynx family, Lypd6 is a glycophosphatidylinositol (GPI)-bound membrane protein known to positively modulate nicotinic cholinergic signaling (Darvas et al., 2009; Miwa et al., 2012). Lypd6 may increase SST interneuron activity by impacting postsynaptic nicotinic modulation of excitatory synaptic inputs, such as in the case of hippocampal CA1 O-LM interneurons where nAChRα2 gates LTP through receptor-mediated calcium influx (Jia et al., 2010). Alternatively, Lypd6 may modulate nicotinic signaling at presynaptic SST terminals to facilitate the release of GABA (Marchi and Grilli, 2010). Either potential mechanism can achieve the end goal of facilitation of SST interneuron activity. However, Lypd6 could mediate additional mechanisms that less directly influence nicotinic signaling. To date, the common sub-cellular localization of Lynx family of proteins and mechanism of nicotinic signaling modulation is largely unknown. However, evidence suggests that the modulatory mechanism of Lypd6 may be complex, that it may localize to synaptic compartments to also regulate ligand sensitivity, or assembly and membrane insertion of nAChRs (Ozhan et al., 2013; Nichols et al., 2014; Thomsen et al., 2014; Puddifoot et al., 2015; Arvaniti et al., 2016). Recently, a closely related Ly6 family protein known as Lypd6b was shown to modulate nAChRs through what seems to be counterintuitive actions – that it increases sensitivity for ACh, yet also reduces net current induced by ACh (Ochoa et al., 2016). Therefore, more mechanistic insight into the physiological action of Lypd6 on nicotinic signaling is eagerly awaited in future studies using new tools for selective targeting and manipulation.

Our study provides circuit level insights into how selective modulation of SST interneurons can trigger rapid disinhibition for enhancing cortical plasticity in adulthood. We demonstrated that the overexpression of Lypd6 in SST interneurons rapidly leads to an experience-dependent increase in their visually evoked activity, and simultaneously a reduction in the visually evoked activity of PV interneurons and an increase in that of excitatory neurons. The suppression of PV interneuron activity and disinhibition of excitatory neurons are typically observed in juvenile mice after the first day of visual deprivation, and can trigger experience-dependent visual plasticity in adulthood when mimicked through chemogenetics (Kuhlman et al., 2013). To our knowledge, this is the first evidence that implicates the role of the SST-PV disinhibitory microcircuit in the adult cortex in enhancing cortical plasticity. Our findings suggest that an experience-dependent increase in SST interneuron activity not only impacts the spatiotemporal coordination of dendritic activity of pyramidal neurons (Silberberg and Markram, 2007; Cichon and Gan, 2015), but also increase the activity of pyramidal neurons potentially through suppression of PV interneurons. During the critical period, V1 SST interneurons have elevated sensitivity towards cholinergic signaling, which allows sensory experience and arousal to drive plasticity mechanisms including branch-specific dendritic activation and somatic disinhibition of excitatory neurons (Yaeger et al., 2019). The Lypd6 may perhaps be a key to re-engaging cholinergic sensitivity in the SST interneurons of the adult cortex and restoring their circuit function for mediating robust and global levels of experience-dependent plasticity.

While overexpression of Lypd6 in SST interneurons re-capitulated one of the hallmarks of juvenile form of V1 plasticity – rapid reduction of the PV interneuron activity and disinhibition of local excitatory neurons after 1 day of deprivation, it remains undetermined whether the subsequent enhanced OD plasticity after 4 days of MD is the juvenile form marked by depression of deprived eye response, as it was in the case of adult mice that underwent chemogenetic suppression of PV interneurons during 1^st^ day of MD (Kuhlman et al 2013). Rather, our manipulation may have accelerated the initiation of the adult form of OD plasticity that typically takes a week or so of MD and only results in the potentiation of the open eye and no change in deprived eye responses. Our approach to acutely measure contralateral biases of individual neurons through electrophysiology was not feasible to distinguish between deprived eye depression and open eye potentiation. Therefore, future investigation is warranted to tease apart these alternative possibilities. Interestingly, a recent study showed that *silencing* SST interneurons during the entirety of 5 days of visual deprivation induces an adult form of OD plasticity, characterized by an elevation of open ipsilateral eye response (Fu et al., 2015). Whether such a manipulation accompanies any form of disinhibition during the initial phase of visual deprivation is currently unknown. Their study and ours collectively highlight the capabilities of SST interneurons in engaging different forms of plasticity, perhaps dependent on the level of SST interneuron activity (reduced vs. enhanced) and the timing of activity change (during the initial phase within 1 day vs. later homeostatic phase after 5 days). We propose a model where Lypd6 may function as a molecular switchboard for SST interneurons to shift modes and activate the SST interneurons to inhibit PV interneurons (Nakauchi et al., 2007; Leao et al., 2012) while disengaging the VIP-SST disinhibitory circuit that relieves inhibitory action of SST interneurons upon the distal dendrites of pyramidal neurons (Chen et al., 2012; van Versendaal et al., 2012; Fu et al., 2015). In this model, both SST-PV and VIP-SST disinhibitory circuits can co-exist to play complementary roles in the modulation of plasticity, depending on the context and brain state.

The capacity for nicotinic modulation by Lypd6 to induce robust experience-dependent cortical plasticity and functional recovery in the adult cortex opens the possibility of targeting this receptor to treat conditions such as amblyopia, stroke, and traumatic brain injury where hope for recovery in later life is low due to diminished plasticity. The selective expression of Lypd6 in deep layer subpopulations of SST interneurons could limit off-target effects (Timmermann et al., 2012; Wang et al., 2015a; Wang et al., 2015b) and make them attractive as novel therapeutic targets, compared to interventions that do not regard subtype or receptor subunit specificity. Novel pharmacological interventions may be combined with behavioral interventions like physical exercise (Kalogeraki et al., 2014), which may further enhance the impact on SST interneuron activity (Polack et al., 2013; Fu et al., 2014; Reimers et al., 2014; Dipoppa et al., 2016; Pakan et al., 2016).

## Data availability

The data that support the findings of this study are available from the corresponding author (H.M.) upon reasonable request.

## Acknowledgments

This work was supported by National Eye Institute R01EY024918, R01EY 026053, R21EY026702 to H.M., F31EY028829 to M.S., National Institute on Drug Abuse T32 DA007135 to M.P.D., National Institute of Mental Health T32MH096678 to M.S., R21MH106919 to H.M., Knights Templar Eye Foundation to H.M., March of Dimes to H.M., Whitehall Foundation to H.M., and Brain and Behavior Research Foundation to H.M. AZ is a member of the Excellence Cluster Immunosensation. We thank Dr. Klas Kullander (Uppsala University), Dr. Yury Garkun, and Dr. Milo R. Smith (Icahn School of Medicine at Mount Sinai) for their comments, Dr. Ming-Hu Han (Icahn School of Medicine at Mount Sinai) for providing technical expertise on optogenetics and Dr. Yasmin Hurd and Dr. Michael Michaelides (Icahn School of Medicine at Mount Sinai) for their expertise on chemogenetics.

The authors declare no conflicts of interest.

## Author Contributions

M.S., M.P.D., and H.M. designed research; M.S., M.P.D, P.N.B., P.Y., and H.M. performed research; A.Z. contributed reagents and provided expertise; M.S., M.P.D, P.N.B., P.Y., and H.M. analyzed data; M.S., M.P.D, and H.M. wrote the manuscript with assistance from all authors.

